# Impact of Triplet State Population on GFP-type Fluorescence and Photobleaching

**DOI:** 10.1101/2024.02.19.580967

**Authors:** Martin Byrdin, Svetlana Byrdina

## Abstract

Based on recently published parameters (Rane et al. 2023, JPCB 127, 5046-5054) for (rs)EGFP triplet state formation and decay rates and yields, we consider the power density dependence of triplet state population dynamics and its consequences for the application of green fluorescent proteins in biological single molecule fluorescence microscopy. We find that under certain conditions, the photon budget of GFP type fluorescent proteins can be linearly dependent on power density and we propose a possible explanation for such a non-Hirschfeld photobleaching behavior. Moreover, illumination with ms pulses at sub-kHz rates is shown to improve photostability. We stipulate that a judicious choice of excitation wavelength should take into account the triplet state absorption spectrum along with the singlet state absorption spectrum. Formulas are given for the estimation of the effects of such choice as function of the experimental parameters.

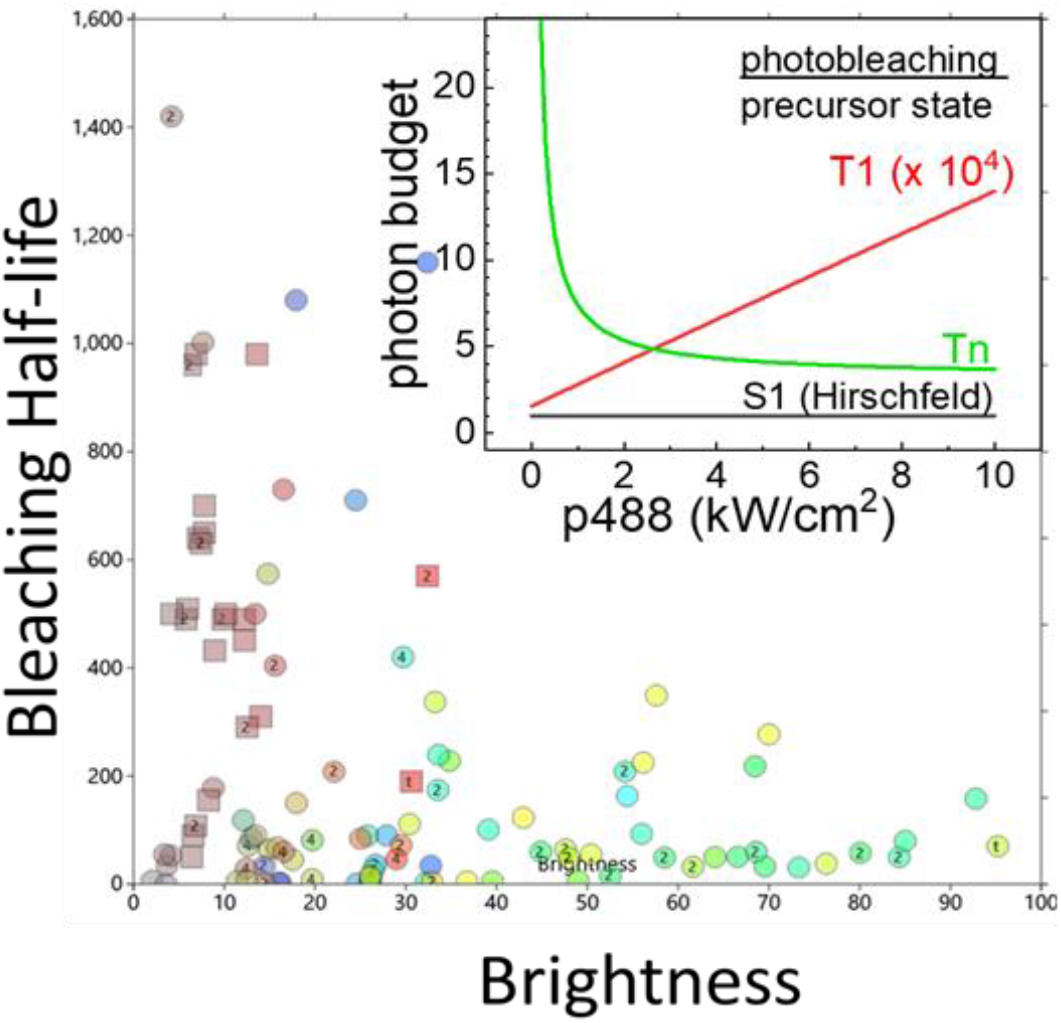

Hirschfeld predicted that photobleaching rates should scale with fluorescence brightness, which is obviously not the case for fluorescent proteins found in the fpbase.org. We investigate a modified theory considering the triplet state as precursor for photobleaching.

## INTRODUCTION

Most microscopists dealing with fluorescent proteins (FPs) have been confronted with two practical inconveniences: undesired premature photobleaching, and the so-called first image artefact.

For many years, huge efforts have been dedicated to the development of new FP variants with decent photostability without sacrificing bright fluorescence. Although it has been recognized since Hirschfeld’s time (Hirschfeld, 1976) that a fluorophore’s molecular brightness is the common factor that scales both the rates of fluorescence and photobleaching, the ancient dream of finding FPs that would be both bright and photostable at the same time has not yet lost its attraction (Urbančič et al., 2013; Wüstner et al., 2014).

On a shorter timescale, the first image artefact (Bierbuesse et al., 2022; Bourges et al., 2023) consists of sometimes severe overexposure upon first switching on laser illumination. Besides being an annoyance for intensity adjustment, this also causes problems for the correct recording of photobleaching time traces. Indeed, with photobleaching often being composed of a reversible fast and an irreversible slower phase, failure to establish a reliable “initial” fluorescence level can lead to largely over-or under-estimating photobleaching halflifes.

Partly as a result of this, experimental photobleaching values are notoriously hard to compare. As an exception, the landmark study of Cranfill and colleagues (Cranfill et al., 2016) used well-defined conditions, protocols and apparatus. Their results for 47 FPs of all colours are plotted in Figure 1 in a “Hirschfeld” presentation showing the inverse photobleaching halflife (i.e. the observed bleaching rate) vs molecular brightness. No clear interdependence seems apparent, not even for FPs within the same colour class.

**Figure 1.**
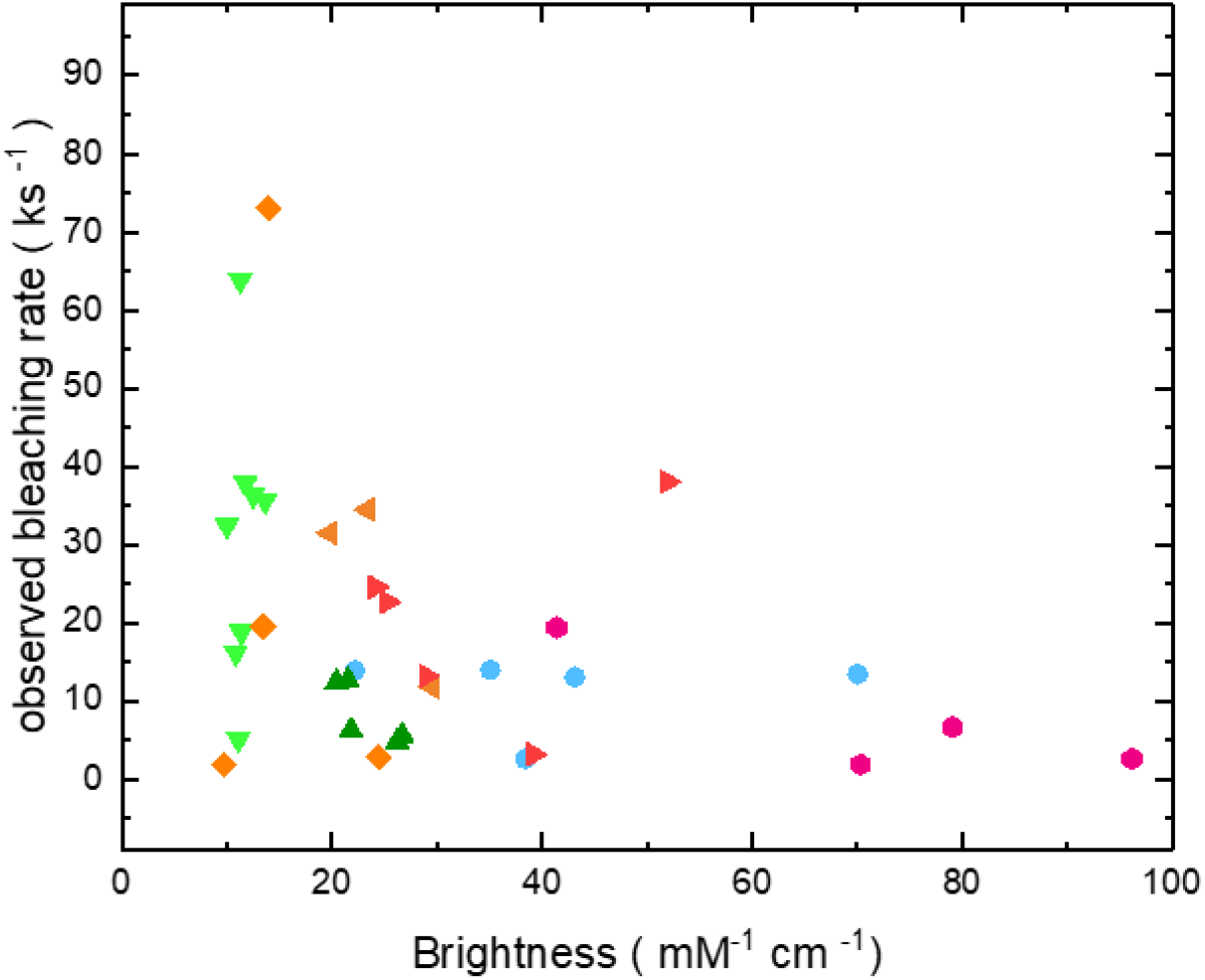
Best available data (for 47 FPs from the landmark paper Cranfill et al.) of bleaching rate vs molecular brightness. Colour coding according to fluorescence colour, same as in tables S1 and S2 in (Cranfill et al., 2016)

Here, we propose a theoretical framework that is based on recent experimental findings on the photophysical properties of the GFP chromophore’s triplet state T1. The correct inclusion of T1 population kinetics into the Hirschfeld approach is proposing possible solutions to both of the abovementioned problems (brightness/photostability trade-off and first image artefact).

Building upon Hirschfeld’s expression for the overall photon budget (PB, the ratio of the intrinsic rate constants for fluorescence and photobleaching *k*_*fl*_ and *k*_*bl*_), we propose a generalization for settings that involve shelved states such as T1, and hence may show non-exponential photobleaching kinetics. Instead of taking the time integral over the complete photobleaching trace (that in most experimental settings is never reached), we consider the ratio G of the instantaneous rates of fluorescence emission and irreversible photobleaching, *k*_*FL*_ and *k*_*BL*_ :

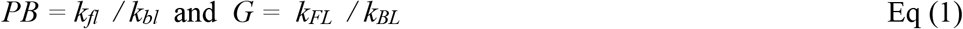

Please note that observable macroscopic rates *k*_*I*_ are denoted by UPPER CASE INDICES whereas intrinsic microscopic rate constants *k*_*i*_ are denoted by lower case indices. Please note also that throughout the paper, we will use normalised populations, leaving out Hirschfeld’s concentration factor *n*. Then, for the classical Hirschfeld case (singlet-only model), our new parameter, called G for “goodness” coincides with Hirschfeld’s photon budget (vide infra), whereas for situations involving reversible dark states such as T1, it contains an excitation power-dependent correction, and will be considered in full detail in this contribution.

The interplay of fluorescence and photobleaching happens at the intersection of several experimental conditions and essential photophysical parameters such as excitation power density (*p*), oxygen availability, illumination timing (continuous *vs* pulsed), temperature,…; and T1 characteristics such as absorption spectrum, lifetime (*τ*_*T*_), quantum yields for its light induced formation via intersystem crossing (ISC, *φ*_*F*_) and depopulation via reverse ISC (RISC, *φ*_*R*_). Most of the latter have previously not been available and therefore not been included in earlier models of FP population dynamics.

In itself, none of the concepts in our model is totally new. Previous work has considered photobleaching starting from T1 or higher triplet states Tn, the role of RISC for population dynamics, use of pulsed excitation to minimize photobleaching, but all of them separately and never “under the same roof”. Here, considering all of these concepts together, we can report for the first time that FP goodness is not excitation power density independent as in the Hirschfeld case, but may increase with increasing excitation power density.

An early triplet photobleaching model by Mondal was hampered by a confusion between irrreversible photobleaching and reversible shelfing, now better known as “blinking” (Mondal, 2008). The rhodamine study by Deschenes et al. considered photobleaching from Tn but did not include RISC (Deschenes & Vanden Bout, 2002). Pulsed excitation schemes as a means to fight photobleaching have been proposed early on, but yielded contradictory results: The fluorescein study by Song et al. (Song et al., 1995) (no RISC considered) and the study by Nishigaki et al. (Nishigaki et al., 2006) on organic dyes found positive effects. In contrast, Wäldchen et al. (Wäldchen et al., 2015), using FPs in cells, found a rather adverse effect, which may be partly due to high laser power and/or the biological reaction of the studied cells. Finally, Boudreau et al. (Boudreau et al., 2016) did a power titration using EGFP in cells under a confocal setting with microsecond pulsing. Apart from this short pulse duration, this experimental study came closest to the conditions considered in the present contribution, and it found 9x reduced photobleaching upon illumination pulsed on the microsecond timescale.

The reminder of this paper is organised as follows: in the THEORY part, we elucidate the key role of T1 population dynamics for fluorescence and photobleaching, before deriving expressions for the power density dependence of the goodness parameter under various illumination conditions. Then, in the EXPERIMENTAL part, we consider paradigmatic rsEGFP2 as an example and apply to it the formulas derived in the theoretical part. We conclude the paper by a DISCUSSION section with focus on practical aspects, such as the role of dissolved oxygen.

## THEORY

### Triplet state buildup and reaction pathways

The important lifetime and formation yield of the triplet state T1 of FPs have for consequence that under the conditions typically found in biological single molecule fluorescence microscopy (immobilized samples under wide-field illumination in the order of kW/cm^2^), significant T1 buildup occurs.

Depending on applied excitation power density, few milliseconds are sufficient to pump the vast majority of the potentially fluorescing molecules into T1 from where they are not only no longer able to fluoresce, causing the well known first image artefact”(Bierbuesse et al., 2022; Bourges et al., 2023). On the other hand, due to the high reactivity of the T1 state, it is also prone to oxidation and reduction reactions which, if not reversed rapidly, may lead to accelerated photobleaching. Increased T1 population thus appears as a double penalty: less fluorescence and more permanent losses. In this contribution, we discuss possibilities and strategies to optimize conditions for fluorescence microscopy to get out of GFP markers a maximum of signal with a minimum of degradation, both for room temperature (RT) and cryo temperature (100 K, CT), and for continuous (cw) and pulsed illumination.

As a starting point, let us consider the photophysical Scheme 1 of electronic states of a GFP-type FP.

**Scheme 1.**
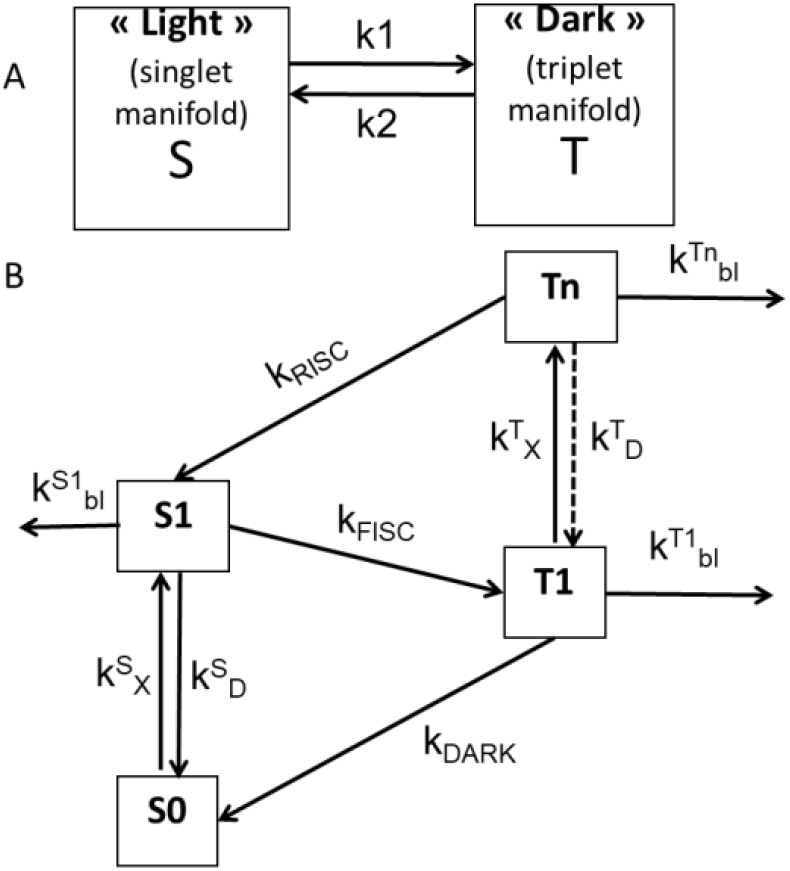
Electronic states and rates involved in FP fluorescence and photobleaching. A, basic two state scheme, B, elaborate four state scheme. Excitation with rate k^S^_X_ pumps molecules from the singlet ground state S0 to the fluorescing excited state S1, from where de-excitation (radiative and non-radiative) occurs with rate k^S^_D_ . Alternatively, forward intersystem crossing (FISC) can populate with rate k_FISC_ the lower lying lowest triplet state T1, from where either dark relaxation with rate constant k_DARK_ repopulates S0 or re-excitation with rate k ^T^_X_ leads to a higher triplet state Tn, from where again de-excitation with rate k^T^_D_ to T1 is possible or reverse intersystem crossing (RISC) with rate k_RISC_ to S1. In parallel to this S/T cycling, all excited states can undergo irreversible photochemistry with rate constants k^i^_bl_.

The relative populations of the four relevant states S0, S1, T1, and Tn are given by the rates connecting them and these in turn are defined by the molecular parameters of the FP chromophore such as absorption cross sections σ_i_, radiative lifetimes τ_i_ and quantum yields φ_i_.

Under continuous illumination, an equilibrium distribution of the FP molecules over the four relevant states S0, S1, T1, and Tn will establish as follows: For a total chromophore population normalized to unity, the final populations for the singlet (S) and triplet (T) manifolds will be given by equation (2):

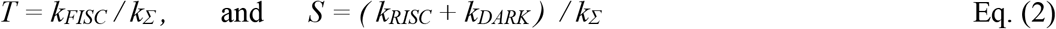

After establishment of equilibrium populations within few milliseconds, we will have for the population ratio S/T shown in Figure 2:

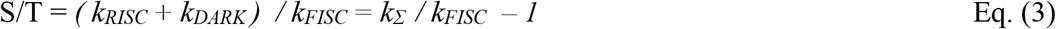

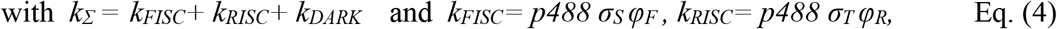

where p488 is the wavelength dependent photon flux density in hc/λ units, *φ*_*F*_ and *φ*_*R*_ are the quantum yields for forward and reverse inter system crossing, *σ*_*S*_ and *σ*_*T*_ are the absorption cross sections at the excitation wavelength of the singlet S0 and triplet T1 states, respectively.

**Figure 2.**
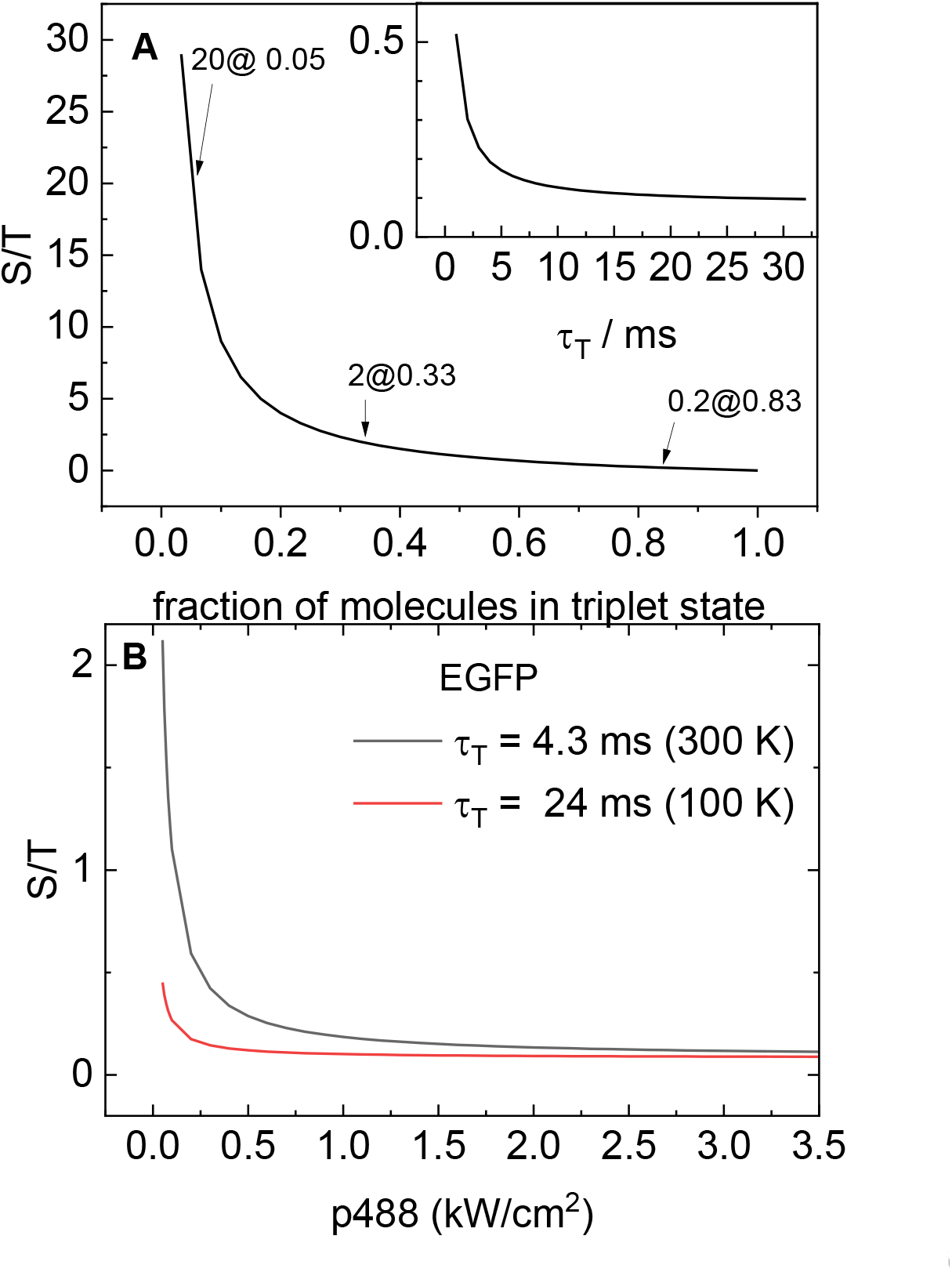
The equilibrium singlet-to-triplet population ratio (eq. 3) decreases rapidly with increased triplet fraction (A, main), increased triplet state lifetime (A inset, for p488 = 1 kW/cm ^2^ and ε_S_ = 50 mM ^-1^ cm^-1^) and increased pump power density p488 (B).

Within each of the S and T manifolds, neglecting for the moment ISC, the fractional population of the excited states is given by the balance of population and depopulation rates:

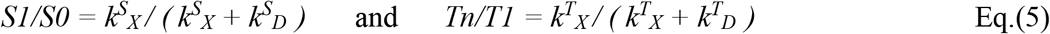

The de-excitation rate of the S1 state *k*^*S*^_*D*_ *= 1/* τ_fl_ *= k*_*fl*_ *+k*_*nr*_ is dominated by fluorescence, that of the Tn state by internal relaxation: *k*^*T*^_*D*_ *= k*_*ir*_ , whereas the T1 dark relaxation rate contains contributions from phosphorescence and quenching: *k*_*DARK*_ *= 1/ τ*_*ph*_ *= k*_*ph*_ *+ k*_*Q*_ .

The excitation rates are given by the product of the applied power density *p488* and the cross section *σ* of the absorbing states:

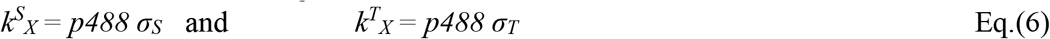

Thus, the behavior of the system is fully defined by the chromophore’s photophysical characteristics and the experimental conditions, mainly excitation light intensity and wavelength, but also temperature, viscosity, quencher concentration, etc. Throughout this paper, and where not mentioned otherwise, we will consider conditions typically found in biological single molecule localization fluorescence microscopy, namely monochromatic excitation at 488 nm with 1 kW/cm ^2^ into the absorption band of the deprotonated EGFP chromophore with ε_S_ = 50 mM ^-1^ cm^-1^. For such excitation conditions, *k*^*S*^_*X*_ ≈ *0*.*8 MHz* which means that for a microscope with a few percent detection efficiency, a per-molecule count rate of few kHz could be expected in the absence of ISC.

Figure 2 shows that with linearly increasing T1 fraction, the photo-physically relevant S/T ratio diminishes hyperbolically, i.e. much faster than linearly.

From equations (4) and (6), it can be seen that all four of the rates *k*^*S*^_*X*_, *k*^*T*^_*X*_, *k*_*FISC*_, and *k*_*RISC*_ are power density dependent. From equations (2) and (5), it follows that so are the relative populations of all four relevant states S0, S1, T1, and Tn. Figure 3 shows the dependence on power density of these populations. Note that for cw power densities going from 10 W/cm^2^ to 10 kW/cm^2^, the equilibrium triplet population increases from 9 to 91%. This is the power density window considered in this paper and illustrated in most of the following figures.

**Figure 3.**
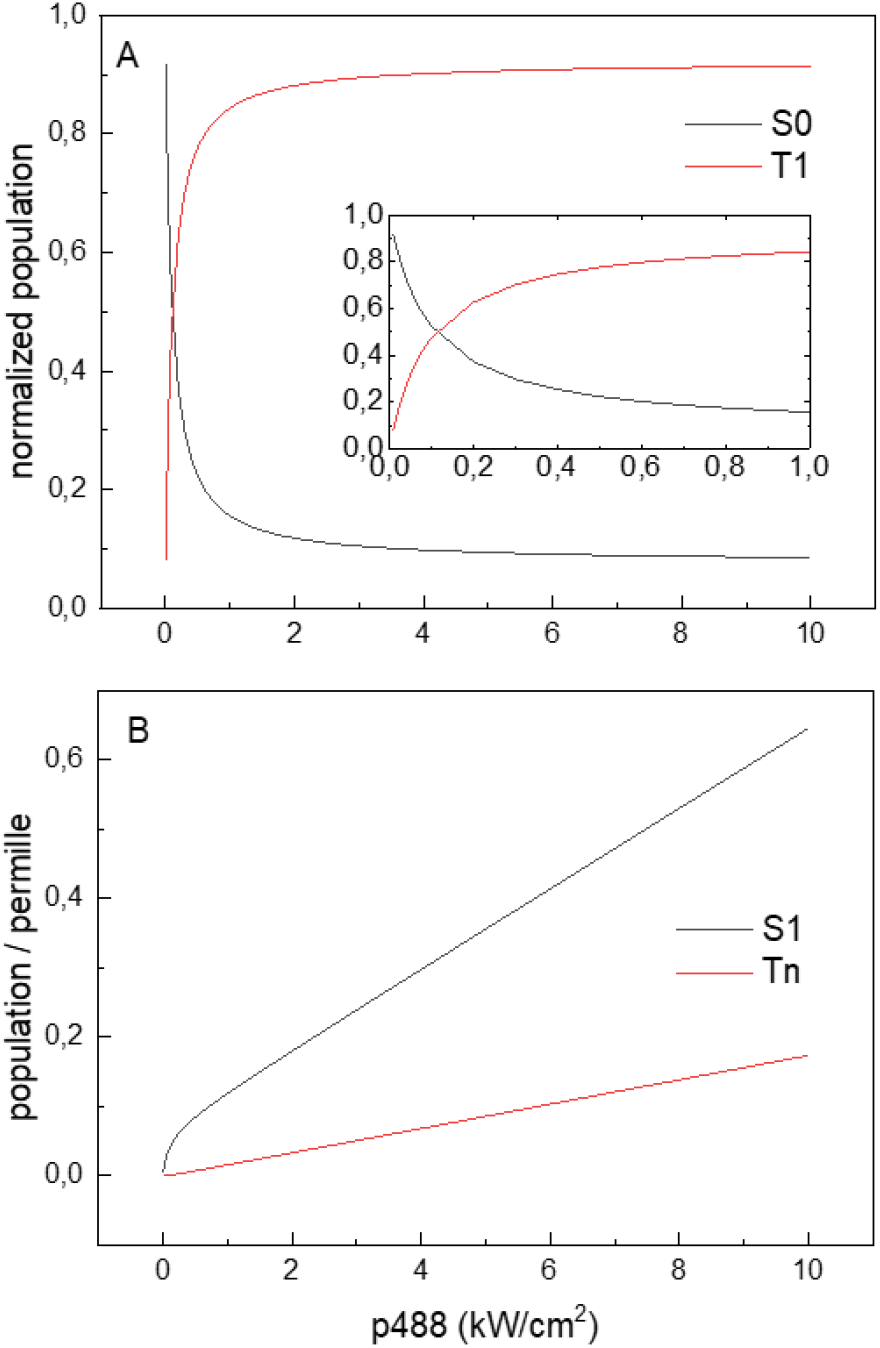
Dependence of equilibrium populations of lowest (A, eq.2) and of higher (B, eq. 5) singlet (black) and triplet (red) states on the applied excitation power density. B, excited state populations, i.e. ground state populations from panel A multiplied by excited state fractions according to eq. (5). The Tn curve is calculated with k_ir_ = 10^10^ Hz and k_DARK_ = 233 Hz, i.e. for RT. Note that the x-scale is common but y-scales differ by 1000 for lower (A) and higher (B) state population panels.

From Figure 3B and equation (5), it can be seen that for any excitation energies < 100 kW/cm^2^, the excited state fractions are less than 1%, so that in the following, we will use S for S0 and T for T1 if not mentioned otherwise.

From equations (2) - (6) it also follows that *Tn* depends on *p488* both via *k*_*FISC/RISC*_ and *via k*^*T*^_*X*_

Thus, in general, an quadratic dependence on power density could be expected.

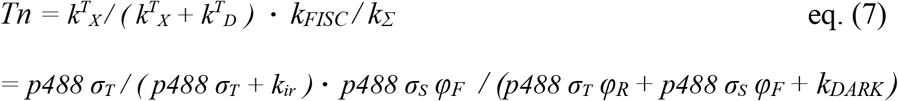

However, Figure 3B shows that the Tn power density dependence appears rather linear. This is due to the fact that in the second term *k*_*DARK*_ contributes little so that p488 cancels out, and in the first term, *k*_*ir*_ dominates the denominator and masks the influence of *p488*. Thus the only apparent p488 dependence is the first numerator, resulting in the apparent linear power density dependence.

### Importance of reverse inter-system crossing

With the quantum yields for triplet state formation and decay as reported by Rane et al. reproduced in Table 1 (Rane et al., 2023), a major fraction of all potentially fluorescent molecules will be shelved away from the fluorescent manifold within few milliseconds, depending on the applied excitation power density.

**Table 1.** Photophysical parameters for rsEGFP2 at RT and CT under illumination at 488 nm in the on state and at 405 nm in the off state.

| rsEGFP2 | From | On RT | On CT | Off RT | Off CT |
| --- | --- | --- | --- | --- | --- |
| Wavelength [nm] |  | 488 | 488 | 405 | 405 |
| $\varepsilon_S$ [mM<sup>-1</sup> cm<sup>-1</sup>] | This work (Fig. 8),<br>Refs. (Rane et al.,<br>2023), (Adam et al.,<br>2022) | 50 | 40 | 25 | 10 |
| $\phi_F$ | (Rane et al.,<br>2023) | 0.003 | 0.003 | 0.003 | 0.003 |
| $\varepsilon_T$ [mM<sup>-1</sup> cm<sup>-1</sup>] | | 10 | 10 | 1 | 1 |
| $\phi_R$ | | 0.001 | 0.001 | 0.001 | 0.001 |
| $k_{DARK}$ [kHz] | $1/\tau_{ph}$ | 0.233 | 0.042 | 0.233 | 0.042 |

One might ask why such a >90% rapid loss of fluorescence intensity should have gone undetected experimentally so far. We propose that this is probably due to the choice of the detection time windows for transient fluorescence spectroscopy: Ultrafast measurements stop recording before the end level is reached, slower measurements miss the “true” initial level and take a “prebleached” state instead. On the contrary, most microscopists are familiar with the routine discarding of the first image due to overexposition.

The T1 shelving process is, however, counteracted upon by the inverse process of light induced back-pumping (RISC), albeit with reduced brightness (quantum yield x cross section, ca 10% of FISC at 488 nm). It is hence the RISC process that determines the distribution of the available FP molecules into steady state populations of S and T (see eq. 2).

Figure 4 shows typical kinetics of equilibrium establishment with and without taking into account the RISC pathway. For the inset of this graph, a low power density of 0.3 kW/cm^2^ was chosen such that the timescale corresponds to the establishment of the S⟺T equilibrium. At higher power densities, the equilibrium will establish faster and the differences of equilibrium fluorescence between with and without RISC will be more important (main panel). At 1 kW/cm^2^, e.g., the final fluorescence levels will differ by a factor of almost 2 for room temperature (RT) and more than 4 for cryo temperature (CT).

**Figure 4.**
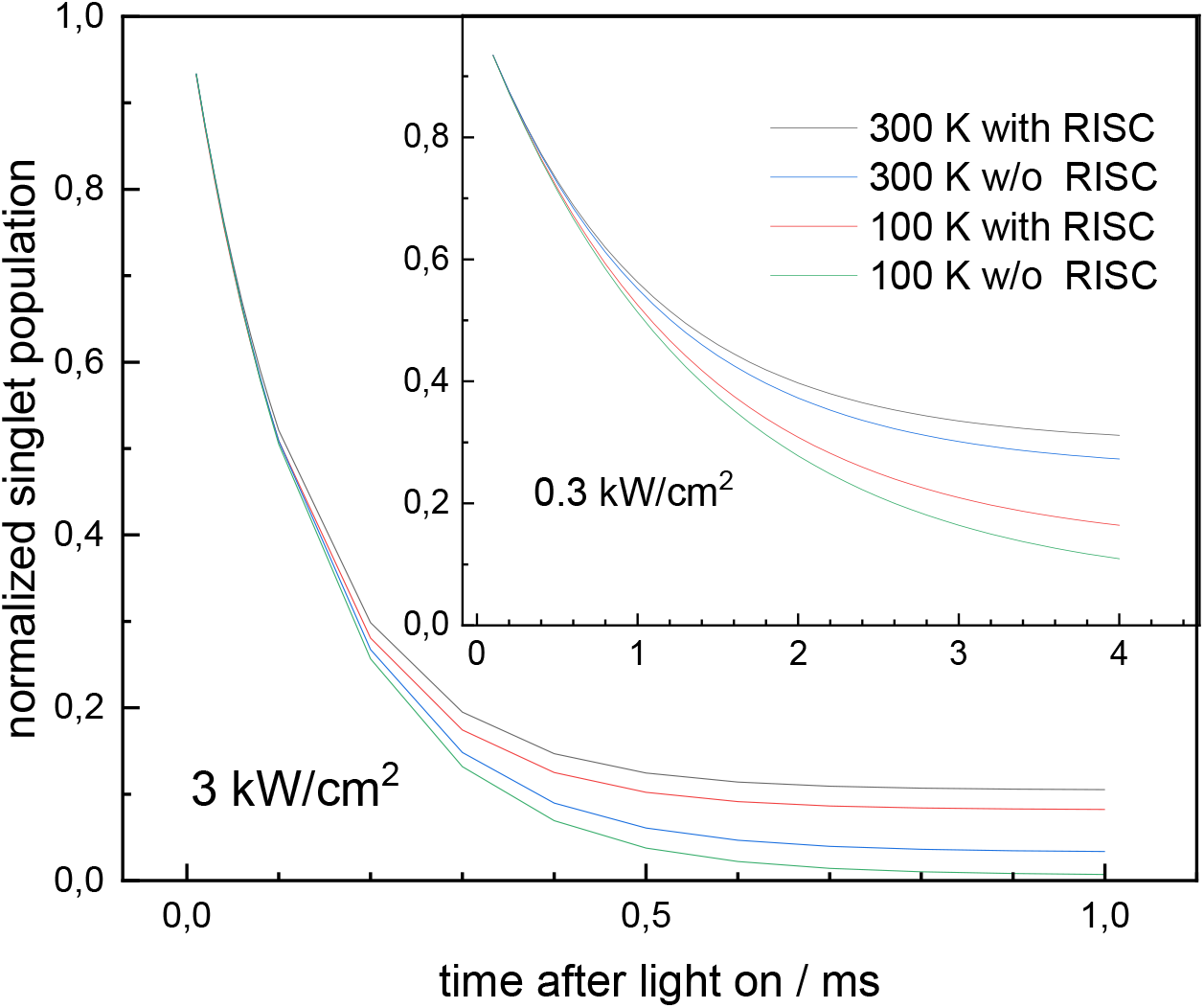
Continuous illumination (here 0.3 kW/cm^2^ in the inset and 3 kW/cm^2^ in the main panel) rapidly pumps most excited fluorophores into the non-fluorescent triplet state. Light-induced reverse intersystem crossing counteracts and partially compensates the loss of fluorescence (black vs blue at RT, red vs green at CT). The kinetics are described by 1-exp (-k_Σ_ t), with the final level being 1-k_FISC_ / k_Σ_.

The incorporation of this additional light induced depopulation pathway of the triplet state complicates the two-state model of scheme 1A (k2 is no longer a purely dark reaction) and necessitates some further elaboration. We will address consequences for fluorescence and photobleaching in the following two sections, separately first for continuous wave illumination and then for illumination by submillisecond pulses with sub-kHz repetition rates. The obtained results will be illustrated on the example of the popular reversibly switchable FP rsEGFP2 in the EXPERIMENTAL section.

Let us define a parameter G (for “goodness”) that describes the two essential properties of a FP: on the up side: the fluorescence FL that can be harnessed from it, and on the down side the photobleaching BL that restricts its practical applicability.

The instantaneous rates of fluorescence emission and irreversible photobleaching, *k*_*FL*_ and *k*_*BL*_ are given by the product of their respective intrinsic rate constants (lower case indices) *k*_*fl*_, *k*_*bl*_ and by the population of the precursor state, which in the Hirschfeld framework is in both cases S1:

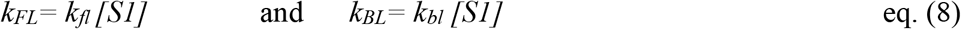

From where it immediately follows that:

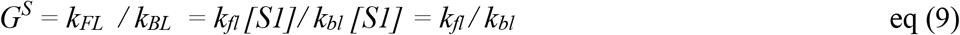

Using the definitions *φ*_*Fl*_ *= k*_*fl*_ */( k*_*fl*_ *+ k*_*bl*_ *+ k*_*nr*_) and *φ*_*Bl*_ *= k*_*bl*_ */( k*_*fl*_ *+ k*_*bl*_ *+ k*_*nr*_*), k*_*nr*_ being the FP’s nonradiative decay rate constant, eq (9) can also be written as:

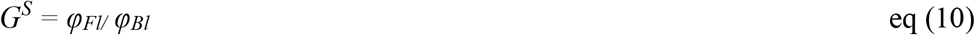

i.e. in the singlet-only case, our goodness equals Hirschfeld’s photon budget PB. Indeed, and that was the other remarkable finding by Hirschfeld, his total photon budget depends only on the intrinsic rate constants for fluorescence and photobleaching, but it is totally independent on most other molecular and experimental conditions, such as chromophore brightness, and, most importantly, applied power density (eq. 10).

Now we observed in the introduction an apparent *independence* of photobleaching rate on the chromophore’s brightness, indicating that Hirschfeld’s initial assumption of photobleaching starting exclusively from S1 is not generally warranted for FPs. We therefore will now consider two limiting cases:

i) the photobleaching precursor state is exclusively T1, or
ii) the photobleaching precursor state is exclusively Tn.

In these two cases, in contrast to the situation considered by Hirschfeld , where S1 is the only photobleaching precursor state, we expect a dependence of G^T^ on power density via the T population (see Figure 3). Comparing the theoretical dependences on power density predicted here with those observed experimentally, eg by Cranfill et al. (Cranfill et al., 2016), we can then strive to make conclusions on the relative contributions of the respective proposed pathways i) and ii).

### Expressions for Goodness under continuous illumination

Let us start considering the case of constant laser illumination, i.e. no pulsing or scanning, consequently the equilibrium populations established after the first few milliseconds do not change any more and we have steady state conditions.

i) *The photobleaching precursor state is exclusively T1*

With eq (8) and the steady state populations from eq (2) we have

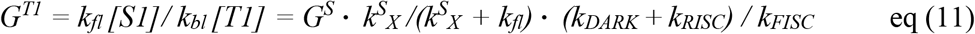

In this equation, the second factor defines the excited fraction of the singlet manifold (eq. 5), and the last factor gives the ratio of the normalized steady state populations of the singlet and triplet manifolds under continuous excitation (eq. 3).To gain insight into the dependence of G^T1^ on the excitation power density *p488* and on the respective singlet and triplet state absorption cross sections *σ*_*S*_ and *σ*_*T*_, we use eq (4) and (6) to explicitly express the rates *k*^*S*^_*X*_ and *k*_*FISC*_ and *k*_*RISC*_ through them. Then we have from eq (11):

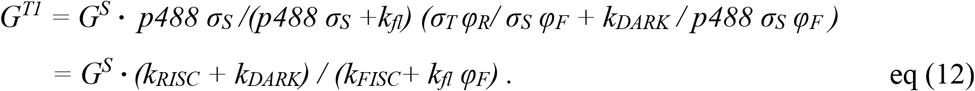

In equation (12), nominally, both *k*_*RISC*_ and *k*_*FISC*_ are dependent on *p488* via the absorption cross sections of their respective precursor states *σ*_*S*_ and *σ*_*T*_. If the other contributions in eq.(12), *viz*. triplet dark recovery in the numerator and fluorescence in the denominator were negligible, *p488* would cancel out and we had a power-independent goodness, just as in the singlet case. However, in fact it is the fluorescence term that dominates the denominator and masks the power dependence of the FISC term. Consequently, in the graphical presentation shown in Fig. 5, we observe rather a linear dependence of G^T1^ on p488. The at first glance counterintuitive observation that the goodness G^T1^ increases with increasing excitation power density is explained by the fact that the RISC pathway effectively limits accumulation of the vulnerable triplet state to the final level of 1-*k*_*RISC*_*/k*_*FISC*_ = 91%, whereas fluorescence continues to increase via enhanced singlet state excitation.

**Figure 5.**
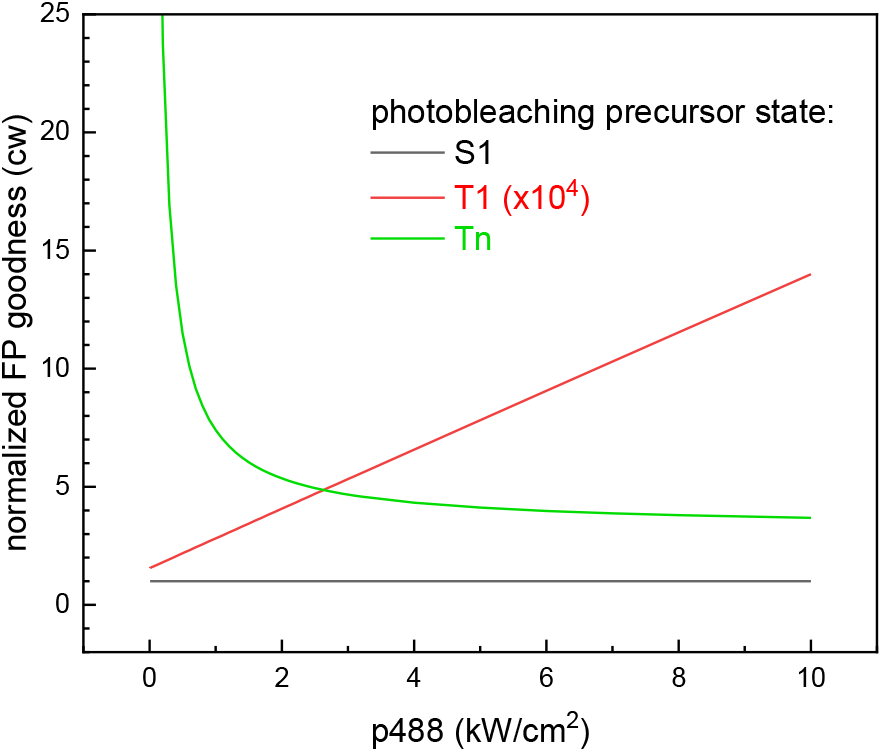
Dependence of Goodness on cw excitation power density for the cases that the precursor state of photobleaching is either S1, T1, or Tn.

ii) *the photobleaching precursor state is exclusively Tn*

With eq (8) we have

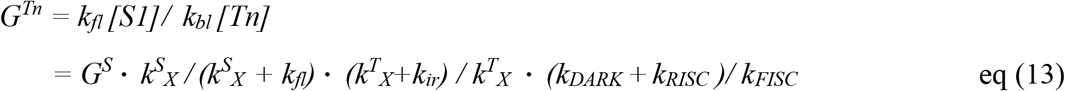

In contrast to eq (11), here we have introduced the additional term *k*^*T*^_*X*_ */ (k*^*T*^_*X*_ *+ k*_*ir*_) for triplet state re-excitation, whereas the overall population factors stay the same as in the previous case (i.e. we assume that the photobleaching via Tn occurs independently from the RISC pathway). Following a completely analogous treatment to that in the T1 case, we obtain the following expression for the goodness:

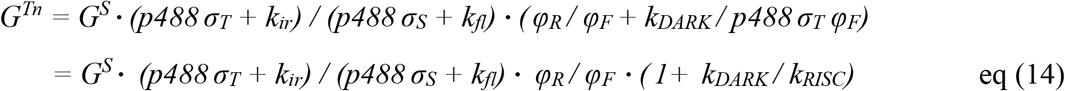

As with eq (7) for [Tn], the excitation power density enters *G*^*Tn*^ twice, both via singlet and triplet state absorption, i.e. under certain circumstances, an inverse quadratic power density dependence can be expected. However, in both instances a second term adds to the power density term and modulates its impact. Again, for not too high excitation, in the first factor, the intrinsic decay rate constants *k*_*fl*_ and also *k*_*ir*_ will outcompete the power density dependent contributions by far because higher triplet states are reputed to be extremely shortlived, in analogy to the Kasha rule for the singlet manifold. As a result, the first factor is roughly ten (*i*.*e. k*_*ir*_ */ k*_*fl*_), and we are left not with an inverse quadratic but rather with an inverse *linear* power density dependence, via *k*_*RISC*_ in the last denominator, in this case. The power density dependences for both triplet states proposed as photobleaching precursors here are shown in Figure 5 together with the Hirschfeld case with S1 as the only precursor.

Note that in Figure 5, the values for a T1 precursor are sub-permille and for better comparison had to be upscaled by a factor of 10000. This difference in scale is due to the fact that [T1] in the denominator is the only “ground” state species, while S1 and Tn are excited and hence populated >1000 times less than their corresponding ground states, see Fig. 3).

To judge the actual impact of the T1 pathway for photobleaching, apart from the population-based goodness presented in Fig. 5, we also need to know the intrinsic rate constant for the bleaching reaction starting from T1, in eq.(11) contained in the *G*^*S*^ term. Such information is currently not available, but it is conceivable that *k*_*bl*_ ^*T1*^ is smaller than those for S1 and Tn, precisely because being a ground state, T1 is lower in energy than S1 and Tn, by 0.76 and 1.37 eV, respectively (Byrdin et al., 2018; Rane et al., 2023).

In conclusion to this section dealing with monochromatic continuous excitation, we relate the theoretically predicted power density dependences of photobleaching (equations (12) and (14), Fig. 5) to those observed experimentally in the most comprehensive study in (Cranfill et al., 2016). There, the authors expected to find a linear power density dependence of the *observed* bleaching rate, i.e. a power density independent goodness. Instead, they found that most FPs showed “supralinear photobleaching” with power exponents scattered between 0.65 and 1.92 with a median around 1.1, their mean ± s.d. is 1.16 ± 0.24. Whereas most of the photobleaching they observed was stronger than linearly power density dependent, some proteins showed sublinear photobleaching. This intriguing behavior might find an explanation if the observed photobleaching rates actually did result from a (partial) compensation of two opposing effects, *viz* inverse linear photobleaching from Tn (case ii, green curve in Fig. 5) and direct linear increase of the goodness from T1 population as predicted for case i (red curve in Fig. 5). In such a scenario, power coefficients scattered around 1.x would not be totally unexpected. To verify these speculations, more experimental work is definitely needed.

### Expressions for Goodness under pulsed excitation

It was proposed early on that potentially harmful exposure of T1 to excitation light could be minimized via a pulsed illumination scheme, e.g. for D-rex or FCS applications, but also for widefield microscopy (Boudreau et al., 2016; Donnert et al., 2007; Gregor et al., 2005; Nishigaki et al., 2006; Ringemann et al., 2008; Song et al., 1995; Wäldchen et al., 2015).

Most of those early attempts traditionally concentrated on MHz pulsing, partly because organic dye triplet lifetimes are found rather in the microsecond range. On the other hand, such timing is natural for MW / cm^2^ excitation power densities where the singlet population decays within sub-microseconds (downscale the x-axes of Fig. 4 by a factor of thousand).

Here, we consider rather kW / cm^2^ excitation power densities and consequently, sub-kHz pulsing schemes with submillisecond exposure times.

To limit accumulation of the triplet state, light exposure time periods should be short with respect to establishment of equilibrium (i.e. submilliseconds) and interleaved with periods allowing for recovery of the singlet population, i.e. few milliseconds at RT and few tens of milliseconds at CT. Under such conditions, no significant triplet state accumulation should occur, but concomitantly the time-averaged fluorescence is reduced due to the dark periods without excitation. In a remarkable 2005 paper, Gregor et al. proposed a formalism to find time-averaged populations for such illumination schemes (Scheme 2) and derived formulas for their calculation. (Gregor et al., 2005)

**Scheme 2.**
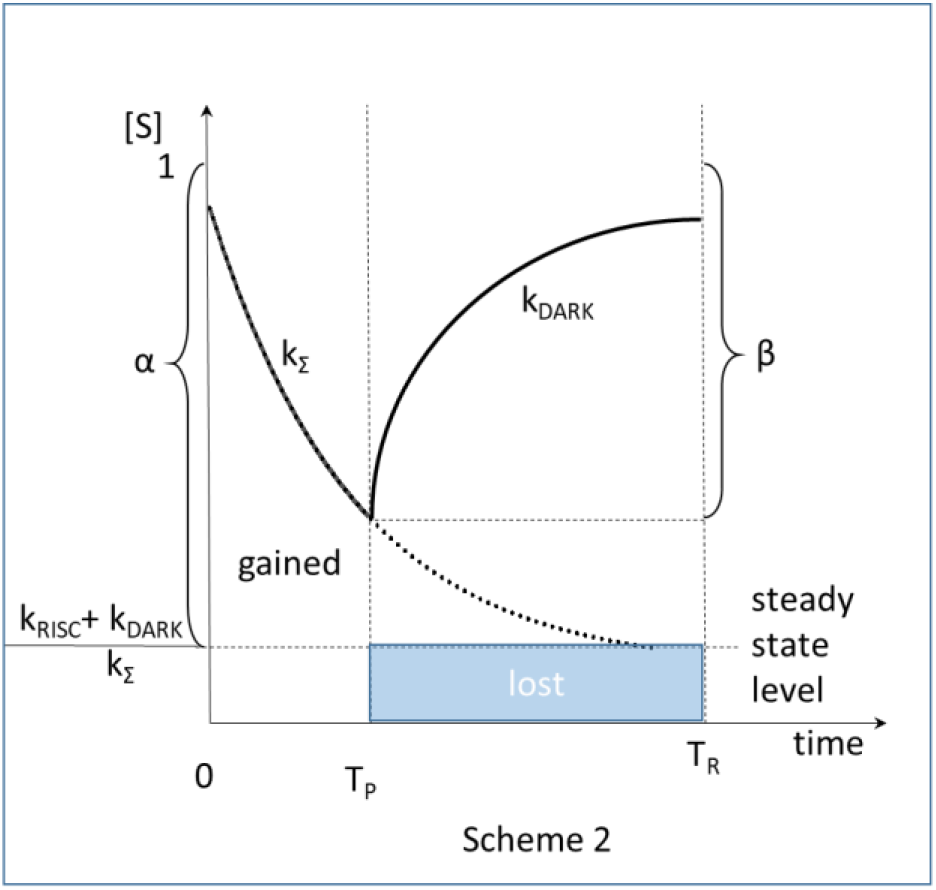
Population evolution under pulsed illumination. The area under the thick black curve is singlet population (i.e. potentially fluorescing), the area above the thick black curve is triplet population (i.e. potentially photobleaching). Under pulsed illumination, singlet state population cycles between a maximal value at the end of a repetition period of duration T_R_ and a minimal value 1-β at the end of a pulse of duration T_P_. As compared to continuous illumination, where at steady state the fluorescence stays at the blue level, during pulses short with respect to 1/ k_Σ_, considerable fluorescence can be gained.

Here, we apply their formalism for the system in Scheme 1B, taking into account the additional RISC pathway, that was not considered in Gregor et al.. Using the obtained expressions, we can generalize the cw treatment of the previous section for the case of interleaved excitation and find optimal parameters for pulse duration *T*_*P*_ and repetition period *T*_*R*_.

For the calculation of *G*_*pulsed*_, we need to obtain expressions for S1 averaged over the time period from zero to *T*_*P*_ (the light period only) for the numerator (fluorescence), and for the denominator (photobleaching) we need T1 averaged over the complete pulse cycle (light and dark, from zero to *T*_*R*_) for *G*^*T1*^_*pulsed*_.

For *G*^*Tn*^_*pulsed*_, we need to average T1 only over the light period from zero to *T*_*P*_, and weight it with the excited fraction *k*^*T*^_*X*_*/(k*^*T*^_*X*_ *+ k*^*T*^_*D*_ ), as in the cw case.

Then we have:

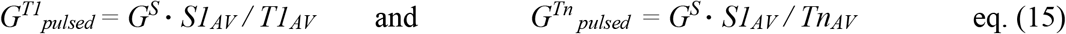

with

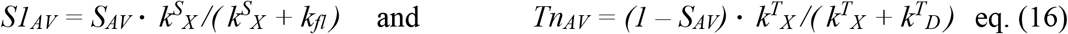

where

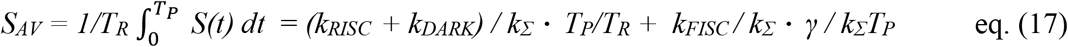

and

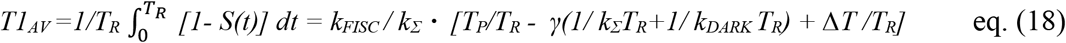

where we introduced *γ* = *[1-exp(-k*_Σ_*T*_*P*_*)][1-exp(-k*_*DARK*_ Δ*T )]/[1-exp(-k*_Σ_*T*_*P*_ *-k*_*DARK*_ Δ*T)]*.with Δ*T= T*_*R*_*-T*_*P*_ being the duration of the dark period between consecutive light pulses. Ffor the detailed derivation of eqs. (17) and (18), see the appendix.

In Eq.(17), the first term describes the “losses” ( i.e. the steady state average value shortened by *T*_*P*_*/T*_*R*_) and the second term describes the “gains”, i.e. additional fluorescence from the not yet depleted singlet state. For the lower excitation energies (i.e. small *k*_Σ_), these gains can be non-negligible, as illustrated in Figure 6A. There exists an optimum for light and dark periods of approximately equal lengths of about 0.7 ms. However, if we consider the *total* fluorescence (Figure 6B), the overall picture is dominated by the “loss” term, whose contribution grows linearly with longer pulses and shrinks with longer recovery periods.

**Figure 6.**
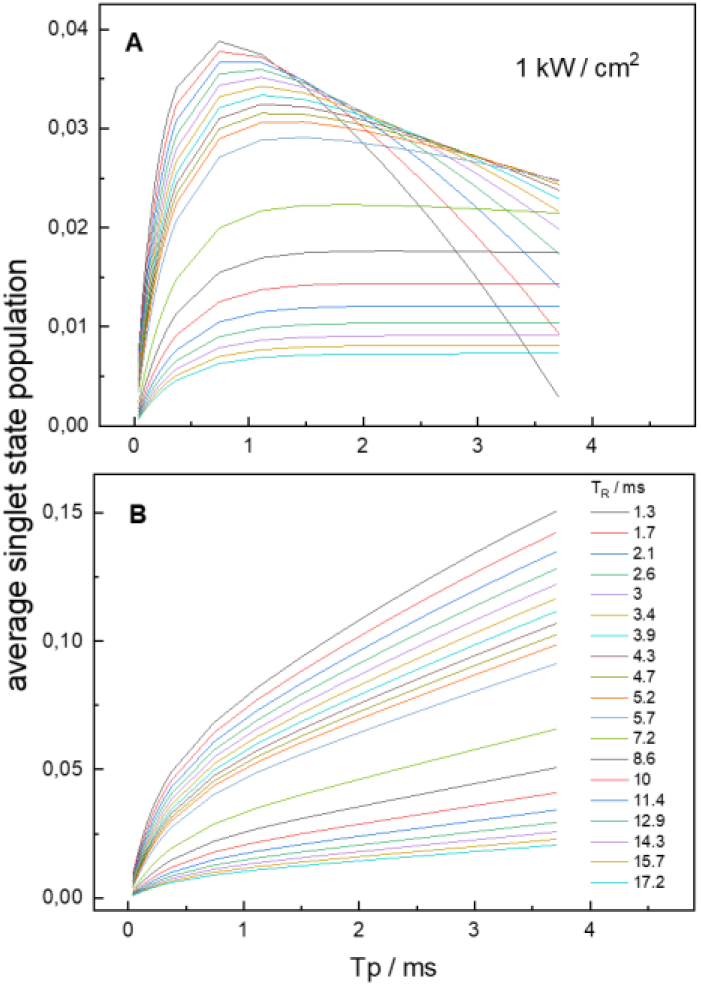
shows the dependence of obtainable fluorescence gain as function of pulse length T_P_ (x axes) and repetition time T_R_ (color coded). As expected, the gain becomes rapidly smaller for longer light pulses and dark periods. A, gain term alone (second term in eq (17)), B total fluorescence (both terms in eq. (17)). Note the differing Y-scales.

Therefore, even under optimal pulse timing, the total fluorescence will never exceed the steady state level under continuous illumination.

Even so, concomitantly with the increased averaged singlet population, less average triplet population accumulates; leading in turn to less photobleaching, and consequently the goodness will be improved for pulsed illumination with respect to continuous illumination.

In eq. (18), the first two terms represent the complement to unity during the light phase (time zero to T_P_) and the second two terms describe the triplet population remaining during dark recovery of the singlet state population (T_P_ to T_R_). To estimate the obtainable gain with respect to the cw situation, we need to replace the total cw population factors for S and T = 1 – S in the cw treatment above (Fig. 3A) with those obtained via equations (17) and (18), while the excited state fractions (Fig. 3B) stay unmodified because establishment of these equilibria S0 ⟺ S1 and T ⟺ Tn is fast (Gigahertz) with respect to millisecond pulsing schemes.

To calculate the power density dependence of the populations under pulsed excitation, we fix *T*_*P*_ and *T*_*R*_ to the optimal values from Fig. 6A as follows:

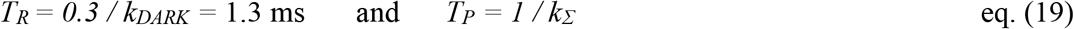

Thus, *T*_*P*_ will be power dependent via *k*_*X*_ entering both *k*_*RISC*_ and *k*_*FISC*_ in *k*_Σ_ . Moreover, explicit power density dependencies will occur via *k*_*RISC*_ and *k*_*FISC*_ and in the exponential expression for *γ*.

In Figure 7, we present populations (A) and goodnesses (B), for pulsed excitation as a function of power density (eqs. 15-18).

**Figure 7.**
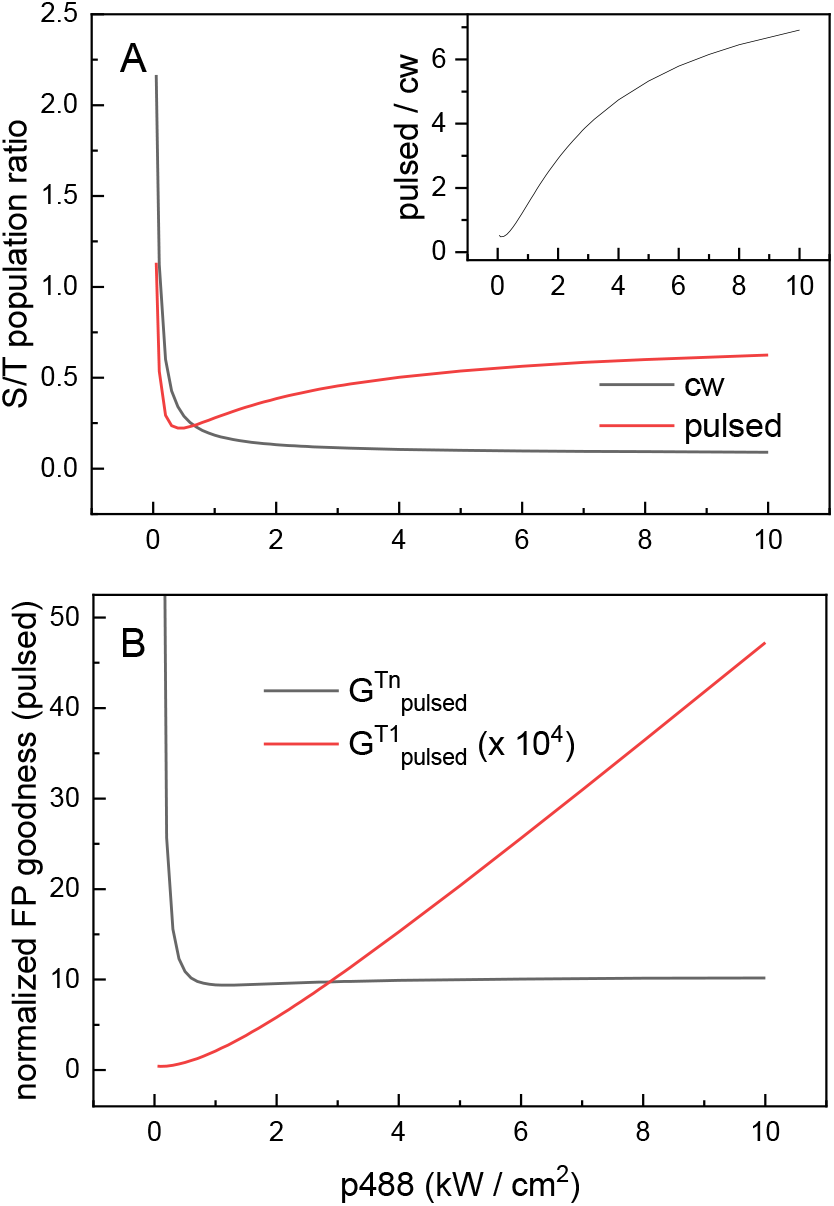
Dependence of singlet-to-triplet manifold populations (eqs.17,18) on excitation power density for continuous (black) and pulsed (red) illumination. Inset, ratio of the two curves in the main panel. A, total populations, B, populations weighted with excited state fractions (equations 15 and 16).

The fact to switch off the excitation light before the singlet population reaches its lowest equilibrium level, allows maintaining the triplet state population at a relatively low level, and for higher excitation even more so. This has for consequence that, in contrast to cw excitation, the S/T ratio for conditions of optimal pulse length (red curve in Fig. 7A) does not decrease but *increases* slightly with excitation power density. This increase is mirrored by the Tn curve in Figure 7B and is in contrast to the cw behavior shown in Fig. 5, despite an overall resemblance between the graphs 5 and 7B. Because the cw equilibrium S/T ratio (black curve in Fig. 7A) decreases with increasing power density, for higher excitation, the pulsed-over-continuous gain is higher (Fig. 7A inset) but levels off at a factor above about six , very close to what was observed experimentally (Boudreau et al., 2016). Interestingly, this gain in effect using pulsed *vs* cw illumination is of the same order of magnitude as the effect observed for *off-*to*-on* photoswitching rsEGFP2 at CT. (Mantovanelli et al., 2023)

## EXPERIMENTAL

### rsEGFP2 *off* state at cryo temperatures also forms a long lived triplet state

For the sake of illustration, in this section we will apply the formulas derived in the THEORY section to a popular reversibly switchable FP, viz. rsEGFP2 at RT and CT for its *on* and also for its *off* state. Indeed, for this protein it was found recently, that its *on* state forms a triplet state the absorption spectrum of which is indistinguishable from that of EGFP. In the same work, we published a straightforward means for the detection of triplet state lifetimes via the kinetics of phosphorescence decay (Rane et al., 2023). For this, for several milliseconds, the sample is exposed to high laser power in order to pump most of the FP molecules into the triplet state. During this period, the photomultiplier is protected by a closed mechanical shutter and a longpass filter. Immediately after switching off the laser, the shutter is opened and the time course of the phosphorescence decay is recorded, yielding transients as those shown in Figure 8B. Using this method, we also investigated a sample of rsEGFP2 that had been exposed at 100 K to 488 nm illumination until it contained predominantly protein in the *off*-state (Figure 8A, red line). Interestingly, after a short pulse of subsequent 405 nm illumination (exciting exclusively the *off*-state), this sample showed also a phosphorescence decay with a lifetime of 24 ms, just as for the protein in the *on* state. Excitation with 488 nm light as a control showed that this signal did not stem from residual *on* state (Fig. 8B). We thus conclude that the *off* state of rsEGFP2 also forms a triplet state with a 100 K lifetime of 24 ms. Due to limitations in signal-to-noise ratio, we were not able to obtain an absorption spectrum for this state and use therefore that of the *on*-triplet as basis for *off*-state calculations in analogy to those performed in the previous sections for the *on* state, just using the absorption coefficients at 405 nm instead of those at 488 nm.

**Figure 8.**
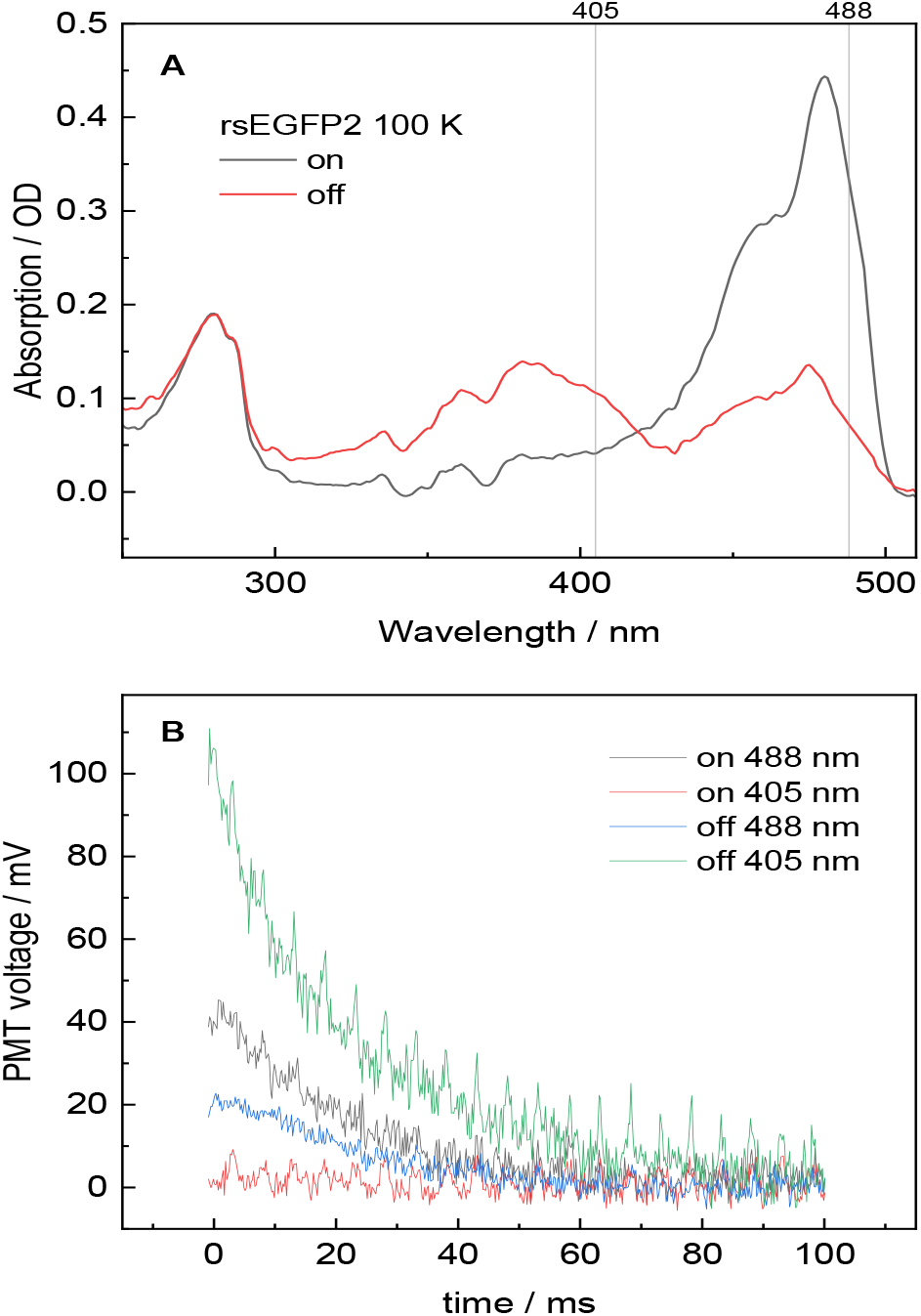
The off-switching of rsEGFP2 reduces 488 nm excited phosphorescence and substantially induces 405 nm excited phosphorescence. A) absorption spectra of a rsEGFP2 drop before (black) and after (red) prolonged 488 nm illumination. B) decay of rsEGFP2 phosphorescence of the two samples in A) after 20 ms of laser illumination with 1 kW cm^-2^ at 488 nm (black and blue) or 405 nm (red and green) light. 256 kinetic traces were averaged. This extensive averaging made appear a 200 Hz correlated noise pattern (especially well visible in the green curve). This noise is present before illumination as well and hence not light induced. In order to show the unmanipulated data, we refrained from applying a digital filter. All curves can be fitted by a monoexponential decay with a time constant of 24 ms suggesting a common precursor state.

### rsEGFP2 goodness for on and off states at RT and CT under continuous and pulsed illumination

Upon the grounds of the previous subsection and based on the photophysical parameters summarized in Table 1, we can investigate the respective influence of *k*_*DARK*_ and of the singlet and triplet state absorption cross sections (405 vs 488 nm) on *k*_*RISC*_ and *k*_*FISC*_. The resulting goodness parameters for both states and both temperatures are summarized in Table 2 for continuous and pulsed illumination, calculated with 1 kW /cm^2^ and in Table 3, calculated for a hundred times lower excitation power density as typically applied in biological widefield microscopy The excitation power density enters the expressions for *k*^*S*^_*X*_, *k*^*T*^_*X*_ , *k*_*FISC*_, *k*_*RISC*_, and, via *k*_Σ_, also *T*_*P*_. Consequently, the calculated goodnesses depend in a complicated manner on *p* and cannot be easily extrapolated from the values in Tables 2 and 3 to other power densities. Therefore, column 2 indicates the formulas to be used for calculation of various *G* values for other power densities. For the *on* state at RT, *G* values for *p488* up to 10 kW /cm^2^ can be read directly from Figs. 5 and 7B for cw and pulsed excitation, respectively. Note also that for the sake of comparability, for pulsed illumination, 1 kW/cm^2^ (Table 2) and 10 W/cm^2^ (Table 3) apply only to the light period,and are not corrected to obtain an unchanged average over the complete cycle, which would result in higher excitation rates during the light period.

**Table 2.** Goodness of rsEGFP2 in the on and off states at RT and CT, calculated from the parameters in Table 1 under illumination of 1 kW /cm^2^ (corresponding to 100 mW on a 100 x 100 µm^2^ spot). Excited state relaxation rate constants k^S^_D_ and k^T^_D_ were set to 10^9^ and 10^10^ Hz respectively. G^S^ was normalized to unity because the value of the intrinsic photobleaching quantum yield is not known precisely. Popular estimates range from 10^-4^ to 10^-6^. All other goodness values were calculated with G^S^=1.This normalization is convenient for straightforward comparison between the different conditions. The actual photon budget, however, is also determined by the fluorescence quantum yield φ_Fl_, that naturally is quite different between on and off states.

| rsEGFP2<br>1 kW/cm<sup>2</sup> | From | On RT | On CT | Off RT | Off CT |
| --- | --- | --- | --- | --- | --- |
| $k_X^S$ [kHz] | Eq. 6 | 761 | 609 | 315 | 126 |
| $k_X^T$ [kHz] | Eq. 6 | 152 | 152 | 12.6 | 12.6 |
| $k_{FISC}$ [kHz] | Eq. 4 | 2.28 | 1.83 | 0.946 | 0.378 |
| $k_{RISC}$ [kHz] | Eq. 4 | 0.152 | 0.152 | 0.013 | 0.013 |
| $k_\Sigma$ [kHz] | Eq. 4 | 2.67 | 2.02 | 1.20 | 0.433 |
| $T_{steady\ state}$ | Eq. 2 | 0.854 | 0.904 | 0.791 | 0.875 |
| $S_{steady\ state}$ | Eq. 2 | 0.146 | 0.096 | 0.209 | 0.125 |
| $S1/S$ (x 10<sup>6</sup>) | Eq. 5 | 0.76 | 0.61 | 0.32 | 0.13 |
| $Tn/T$ (x 10<sup>6</sup>) | Eq. 5 | 15 | 15 | 1.3 | 1.3 |
| $G^S$ | Eq. 9 | 1 | 1 | 1 | 1 |
| $G^{T1}$ (x 10<sup>6</sup>) | Eq. 12 | 3810 | 1900 | 2500 | 540 |
| $G^{Tn}$ | Eq. 14 | 8.54 | 4.24 | 66.2 | 14.3 |
| $T_P$ [ms] | Eq. 19 | 0.37 | 0.49 | 0.83 | 2.3 |
| $T_R$ [ms] | Eq. 19 | 1.3 | 7.2 | 1.3 | 7.2 |
| $S_{AV}$ | Eq. 17 | 0.190 | 0.199 | 0.211 | 0.186 |
| $T_{AV}$ | Eq. 18 | 0,323 | 0.248 | 0503 | 0.343 |
| $G^{T1}_{pulsed}$ (x 10<sup>6</sup>) | Eqs. 15-16 | 446 | 491 | 132 | 68.4 |
| $G^{Tn}_{pulsed}$ | Eqs. 15-16 | 11.7 | 9.97 | 66.8 | 22.8 |

**Table 3.** Same as Table 2, but for hundred times lower excitation power density of 10 W/cm^2^, (corresponding to 1 mW on a 100 x 100 µm^2^ spot).

| rsEGFP2<br>10 W/cm<sup>2</sup> | From | On RT | On CT | Off RT | Off CT |
| --- | --- | --- | --- | --- | --- |
| $k_X^S$ [kHz] | Eq. 6 | 7.61 | 6.09 | 3.15 | 1.26 |
| $k_X^T$ [kHz] | Eq. 6 | 1.52 | 1.52 | 0.126 | 0.126 |
| $k_{FISC}$ [Hz] | Eq. 4 | 22.8 | 18.3 | 9.46 | 3.78 |
| $k_{RISC}$ [Hz] | Eq. 4 | 1.52 | 1.52 | 0.13 | 0.13 |
| $k_\Sigma$ [Hz] | Eq. 4 | 267 | 61.5 | 248 | 45.6 |
| T<sub>steady state</sub> | Eq. 2 | 0.09 | 0.297 | 0.038 | 0.083 |
| S<sub>steady state</sub> | Eq. 2 | 0.91 | 0.703 | 0.962 | 0.917 |
| S1/S (x 10<sup>6</sup>) | Eq. 5 | 7.6 | 6.09 | 3.2 | 1.3 |
| Tn/T (x 10<sup>6</sup>) | Eq. 5 | 0.15 | 0.15 | 0.013 | 0.013 |
| G<sup>S</sup> | Eq. 9 | 1 | 1 | 1 | 1 |
| G<sup>T1</sup> (x 10<sup>6</sup>) | Eq. 12 | 2400 | 430 | 2380 | 420 |
| G<sup>Tn</sup> | Eq. 14 | 524 | 94.6 | 6295 | 1104 |
| T<sub>P</sub> [ms] | T<sub>R</sub> / 2 | 0.65 | 3.6 | 0.65 | 3.6 |
| T<sub>R</sub> [ms] | Eq. 19 | 1.3 | 7.2 | 1.3 | 7.2 |
| S<sub>AV</sub> | Eq. 17 | 0.498 | 0.471 | 0.500 | 0.498 |
| T<sub>AV</sub> | Eq. 18 | 0.0436 | 0.149 | 0.0191 | 0.0416 |
| G<sup>T1</sup><sub>pulsed</sub> (x 10<sup>6</sup>) | Eqs. 15-16 | 86.9 | 19.3 | 82.3 | 15.1 |
| G<sup>Tn</sup><sub>pulsed</sub> | Eqs. 15-16 | 49.6 | 35.6 | 249 | 99.2 |

The numeric values in Tables 2 and 3 are calculated from the formulas in column two using the photophysical parameters in Table 1. They are therefore exact values without error bars. However, the uncertainty of the absorption cross sections can be estimated to be in the order of 10%, which relative error will propagate throughout the calculations and as a consequence, the *G* values listed are also to be taken with such a precision of ±10%.

Comparison of Tables 2 and 3 shows as expected, that all the calculated rates and excited state populations are hundred times lower for 10 W/cm^2^ than for 1 kW/cm^2^. In contrast, populations of singlet and triplet manifolds are inversed: 90% triplet at 1 kW/cm^2^ *vs* 90% singlet at 10 W/cm^2^. As a consequence, intuitively, one could expect for the lower excitation power conditions more fluorescence and less photobleaching, hence higher goodness. This is actually the case for *G*^*Tn*^, but the observed effect is mainly due to the 100x diminished Tn population. As T1 is a ground state, no such effect is observed and the goodnesses for the two excitation conditions can be compared directly. As predicted by Figures 5 and 7, and contrary to what intuition said, we observe an up to 25x *increase* of goodness with higher excitation power, both with continuous and pulsed illumination. The reason for this unexpected behavior has been discussed in the THEORY section in the context of eq. (12).

Comparison of columns 3 with 4 and 5 with 6, respectively, shows that, against expectations and observations (Schwartz et al., 2007), lowering the temperature does not increase but rather decrease the goodness, and this independently on the choice of the precursor state (T1 or Tn) or the illumination scheme (cw or pulsed. This effect is due to the fact that at CT, the triplet state dark lifetime is considerably prolonged and it may speak against the assignment of the triplet state as the only photobleaching precursor state. Such a conclusion could also be drawn from comparing G^T1^ between cw and pulsed excitation schemes that always lead to a reduction in goodness due to the fact that bleaching continues during the dark period if T1 is the precursor state. In contrast, bleaching from Tn occurs only during the reduced light period during pulsing and consequently, *G*^*Tn*^ is always considerably increased for pulsed illumination (20% at RT and even 100% at CT).

The comparison of the *G*^*T1*^ and *G*^*Tn*^ rows invariably shows orders of magnitude better goodness for the latter but this is somewhat artificial due to the extremely short Tn lifetime and hence vanishing population fraction (see line *Tn/T*). As *G*^*T1*^ increases linearly with excitation power density (both for cw, Fig. 5 and pulsed, Fig. 7), applying MW / cm^2^ instead of kW / cm^2^ compensates for this effect and renders the absolute *G* values comparable.

Finally, the fact that between *on* and *off* state excitation (488 nm *vs* 405 nm) both singlet and triplet state absorption differ, but the latter more so, allows for a comparison of the relative impact of these two determinants on the goodness. Naturally, the effect of triplet state absorption is highest on *G*^*Tn*^ , where re-excitation of T1 is required. Here, a reduced *ε*_*T*_ results in largely increased photostability. In view of its practical importance, such an effect definitely deserves experimental verification. In contrast to *G*^*Tn*^, *G*^*T1*^ appears to scale linearly with *ε*_*S*_ but to vary little with *ε*_*T*_. If T1 were to be the dominating photobleaching precursor, this would mean that GFP-type FP goodness could be improved by choosing excitation wavelength near the singlet state absorption maximum, which again is matter for experimental verification.

## DISCUSSION

It has long been known that triplet state population plays a decisive role for photobleaching (Deschenes & Vanden Bout, 2002; Jacques et al., 2008). Here we made use of the recently published photophysical parameters for this state for GFP type FPs to propose strategies to optimize a FP’s goodness (or photon budget) (Rane et al., 2023). These should concentrate on the choice of a suitable wavelength of triplet absorption rather than for singlet absorption that, due to the smaller S1 lifetime and population, is expected to contribute less to photobleaching. A crucial role is played by the light induced back pumping of a fraction of the triplet manifold into the singlet manifold (RISC). This phenomenon prevents the formation of 100% triplet population and complicates kinetic analysis. It has previously been studied for organic dye systems by Reindl and Penzkofer using pico-to-nanosecond fluorescence spectroscopy (Reindl & Penzkofer, 1996a, 1996b) and by Widengren and Seidel (Widengren & Seidel, 2000) using FCS (fluorescence correlation spectroscopy). The timescale of those measurements is considerably shorter and the applied pulse power densities correspondingly higher than considered here. Also using FCS, several groups previously studied GFP-like systems in solution (Haupts et al., 1998; Widengren et al., 1999). However, the accessible time window of that technique is limited by diffusion across the illumination spot to submilliseconds. Hence, the observed reversible photophysical phenomena must be ascribed rather to isomerization and/or (de )protonation events and not to intersystem crossing.

Millisecond studies on a GFP mutant by Chirico et al. also were interpreted in the framework of isomerization/ionization (Chirico et al., 2005). Using phosphorescence decay kinetics, a 4.2 ms lifetime was unequivocally determined for the EGFP triplet state at room temperature (Byrdin et al., 2018). The present work is the first to systematically consider the consequences of this exceptional long lifetime for the FP goodness under conditions typical for biological single molecule localization microscopy.

A key role for triplet state quenching is played by molecular oxygen, as underlined by the observation that in EGFP the T1 lifetime at RT is more than doubled in its absence (Byrdin et al., 2018). In the T1 quenching process, singlet oxygen is formed that in turn attacks vulnerable sulphur containing residues (methionine, cysteine) in the chromophore vicinity leading to irreversible photobleaching (Duan et al., 2013). As a consequence, oxygen plays an ambiguous role both as fluorescence enhancer via triplet quenching and as bleaching agent via the resulting singlet oxygen. We modelled the concentration dependence of this double role for four scenarios and the results are presented in Fig. 9.

**Figure 9.**
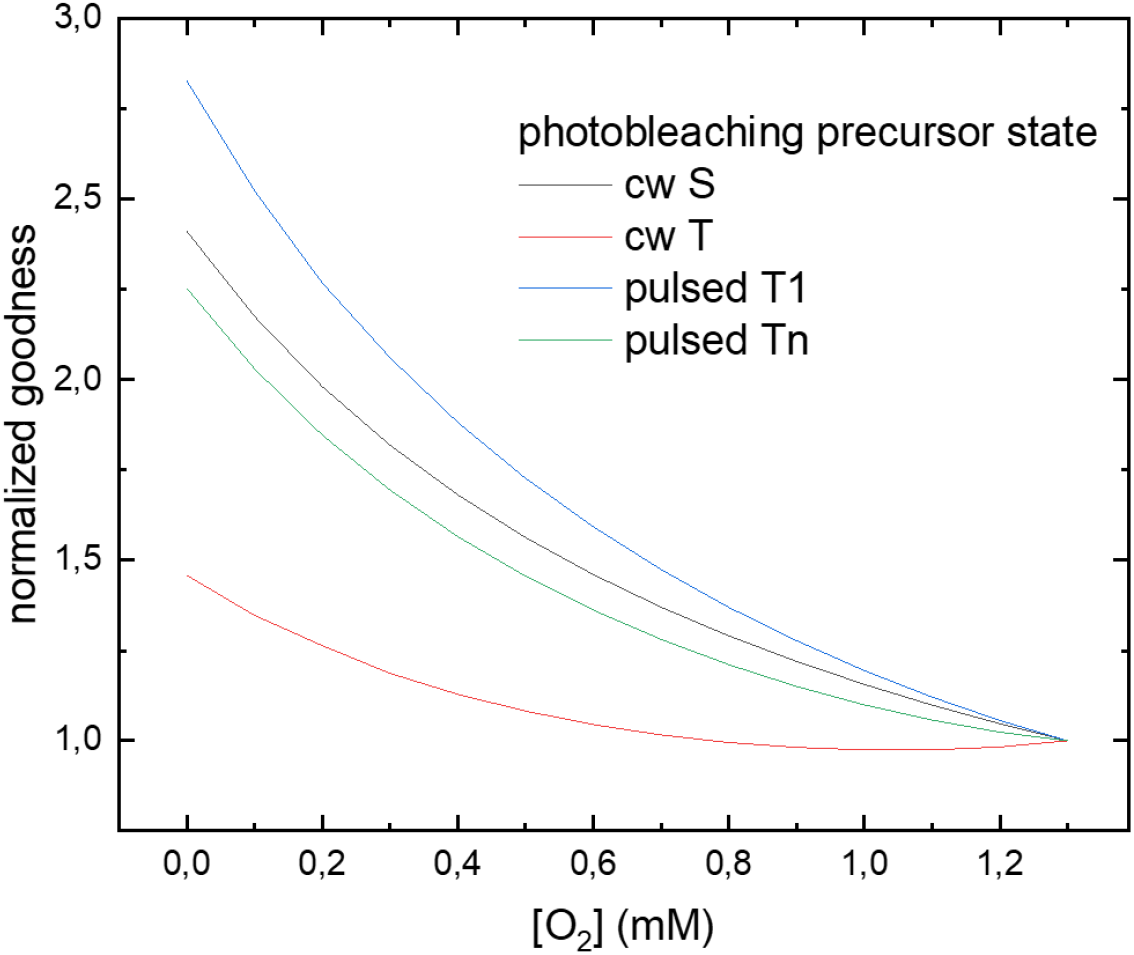
Impact of dissolved oxygen on FP goodness via triplet state quenching and enhanced chromophore vulnerability. The curves are normalized for conditions of oxygen saturation (1.3 mM).

Unsurprisingly, the goodness is best for lowest oxygen concentration in all cases. Interestingly, the oxygen sensitivity of FP goodness is less strong if T is the photobleaching precursor, because in this case the two effects counteract upon each other. Using pulsing schemes reduces the T population and thus returns the system to a S-like behavior with higher sensitivity.

Last but not least, a word is warranted concerning the conditions under which photobleaching is determined experimentally. As discussed, both the excitation power density and the dark decay rate enter the *G*^*T*^*/ G*^*S*^ correction factors. Therefore, both the illumination conditions (power density at sample position) and the sample environment (e.g. temperature, presence of quenchers in the buffer, e.g. oxygen, see above) will affect the observed photobleaching lifetimes and hence the bleaching quantum yield obtained from them. Moreover, a broadband light source can excite singlet and triplet state molecules differently than a monochromatic source. A laser scanning excitation acts as a pulsed source whereas wide-field illumination is continuous and even apparently continuous lasers may by construction actually be high frequency pulsed. The formulas introduced here allow on one hand for the due consideration of such influences, and on the other hand for careful planning of experiments: judicious choice of excitation wavelength, or even use of a second light source for the increase of RISC efficiency, or generally dark state engineering (Ludvikova et al., 2023; Peng et al., 2021).

Concerning presence and concentration of potential quenchers of the triplet state, it should be kept in mind that they may also effect the obtainable fluorescence level via direct interaction with the singlet state, ground or excited (Murray, 2017). Finally, considering the triplet state as potential photobleaching precursor results in non-linear dependence of photobleaching on excitation power density (Figures 5 and 7) and therefore, generally, care is needed in application of the classical Hirschfeld linear photobleaching model for the extraction of numerical photobleaching quantum yield values.

In conclusion,we have shown the enormous importance of triplet state population dynamics for the photophysical behavior of GFP type proteins at RT and CT, both for continuous and pulsed illumination. We found that consideration of the triplet population, yields the goodness of FPs sometimes almost independent of singlet state absorption but linearly dependent on both excitation power density and RISC brightness, with the y-offset being determined by the triplet dark decay rate. The orders of magnitude higher apparent bleaching quantum yield of FPs as compared to organic dyes such as rhodamines is thus explained by the long triplet lifetime of the former, having for consequence a higher steady state triplet population. Use of sub-millisecond excitation pulses with sub-kHz repetition rates is proposed to remedy these effects to a certain extent. Formulas are derived that allow to simulate the fluorescence and photobleaching outcome for various conditions as function of molecular parameters such as fluorescence lifetime, triplet formation and decay quantum yields and lifetimes, as well as pulse timing (duration, frequency). Further experimental investigation of the proposed photobleaching pathways is crucially necessary but even before that will be accomplished, the possibility of photobleaching via the triplet state should be seriously taken into account for the planning of biological fluorescence microscopy experiments, especially at power densities as routinely used for single molecule localisation.

## Acknowledgements

We thank Ninon Zala (Grenoble) for the skillful preparation of the protein samples. The IBS acknowledges integration into the Interdisciplinary Research Institute of Grenoble (IRIG, CEA).

## Appendix

Derivation of eqs. (17, 18) for averaged populations under pulsed illumination. Following the approach by Gregor et al. (Gregor et al., 2005), we consider the simplest possible two state model (Scheme 1A). Under illumination, we have:

*k1 = k*_*FISC*_ and *k2 = k*_*RISC*_ *+ k*_*DARK*_, while in obscurity we will have *k1 = 0* and *k2 = k*_*DARK*_. For normalized total population S + T = 1, under these conditions the total singlet population S will evolve as follows for light (eq. 20) and dark (eq. 21) conditions:

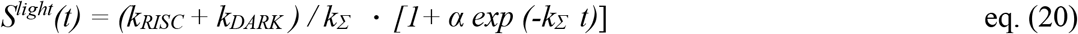

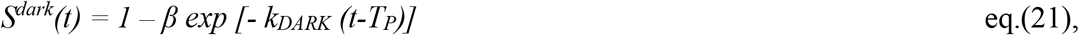

where α and β are amplitude constants that can be determined from equating the populations of eqs (20) and (21) at time points *t = T*_*P*_ (eq. 22) and *t = 0 = T*_*R*_ (eq. 23) (see Scheme 2):

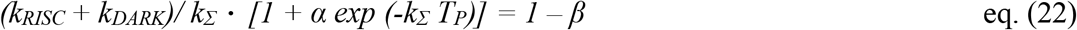

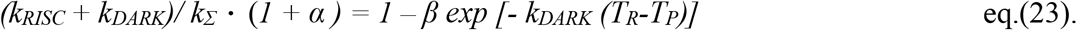

Upon rearrangement, we obtain:

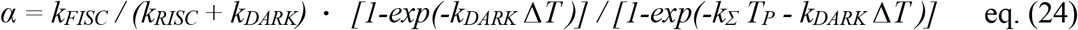

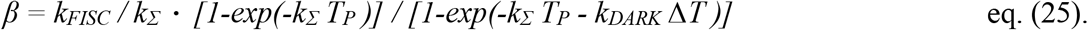

with Δ*T= T*_*R*_ *-T*_*P*_ and *k*_Σ_ *= k*_*FISC*_ *+ k*_*RISC*_ *+ k*_*DARK*_ . The time average of the singlet population contributing to fluorescence during one complete light/dark cycle is then given by eq. (26), where the integration over the S population is done exclusively over the light period from t=0 to t = T_P_. This is because no excitation light is present during the dark recovery period, hence no fluorescence is excited even if potentially fluorescent singlet state molecules are available.

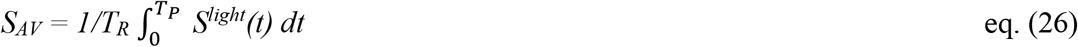

Substituting equations (20) and (24) into eq. (26) and performing the integration yields eq. (17) for the average fluorescing singlet population.

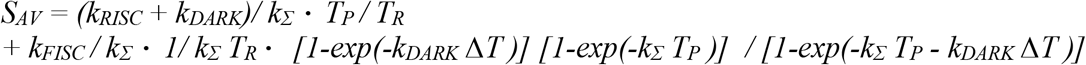

The complement to unity will give the fraction of the triplet population that is prone to photobleaching during the light period. In view of the fact that in case of a T1 precursor state for photobleaching, no further excitation is needed, the average triplet population needs to be calculated including the singlet recovery during the dark period T_P_ toT_R_, yielding eq. (27):

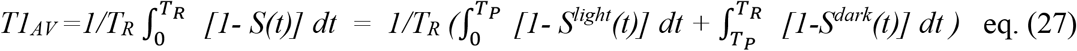

Substituting equations (20),(21), (24) and (25) into eq. (27) and performing the integration yields eq. (18) for the triplet population averaged over one complete light/dark cycle.

